# Spatial profiling of chromatin accessibility reveals alteration of glial cells in Alzheimer’s disease mouse brain

**DOI:** 10.1101/2025.05.01.651759

**Authors:** Dehui Kong, Hongjie He, Joseph Lechner, Jing Yang, Aijun Zhang, Ling Li, Ramkumar Mathur, Diane C. Darland, Motoki Takaku, Lu Lu, Junmin Peng, Kai Yang, Xusheng Wang

**Author notes:** Correspondence authors (K.Y.), (X.W.). These authors contributed equally (Dehui Kong, Hongjie He).

## Abstract

Abnormal epigenetic modifications, including altered chromatin accessibility, have been implicated in the development and progression of Alzheimer’s disease (AD). In this study, we applied spatially resolved chromatin accessibility profiling by performing spatial assay for transposase-accessible chromatin using sequencing (ATAC-seq) to analyze brain tissues from 5×FAD AD model and C57BL/6J control mice. Our analysis identified seven major cell types across 11 brain regions and further characterized glial subtypes—microglia and astrocytes— revealing subtype-specific chromatin accessibility changes in 5×FAD mice relative to controls. These alterations were associated with AD-related pathways, including neuroinflammation and immune dysregulation, and synaptic dysfunction and neuronal signaling in microglia, as well as lysosomal and proteasomal activity, lipoprotein metabolism, and mitochondrial dysfunction in astrocytes. We also characterized cell-type-specific enrichments of motifs and transcription factors, including enrichment of *Bcl11a* in microglia. Furthermore, we linked chromatin accessibility changes in 5×FAD mice to human AD risk genes, highlighting altered epigenetic signatures in genes such as *Trem2*. Supporting the spatial ATAC-seq findings, flow cytometry validated a selective increase in VISTA (encoded by *Vsir*) protein expression in 5×FAD microglia, supporting the spatial ATAC-seq findings and indicating a shift toward a pro-inflammatory, disease-associated microglial (DAM-like) phenotype. These findings underscore the utility of spatial chromatin accessibility profiling for uncovering brain-region specific cell identities and epigenetic mechanisms underlying AD pathogenesis.

## Introduction

Alzheimer’s disease (AD) is a prevalent and progressive form of dementia, defined by hallmark pathological features such as beta-amyloid (Aβ) plaques and Tau neurofibrillary tangles [1,2]. Over the past decade, genome-wide association studies (GWAS) have identified three causative genes (APP, PSEN1, and PSEN2) [3–6] in early-onset familial cases, and about 40 AD risk genes including APOE, TREM2, CLU, PICALM, ABCA7, BIN1 and ADAM10 [7–11]. A significant portion of common variants in AD risk genes are localized within regulatory elements, frequently corresponding to cell-type-specific regions relevant to the disease [12]. Beyond genetic risk genes, recent research has increasingly appreciated the role of epigenomic alterations in AD [13–15]. Chromatin accessibility, as one of the key epigenomic modifications, determines the ability of regulatory factors to bind DNA and regulate gene expression. As chromatin accessibility patterns are highly specific to each cell type, alterations in these patterns can reveal important shifts in cell-type-specific gene regulation. In AD, changes in chromatin accessibility at bulk brain tissues have been associated with the disease onset and progression [16–18].

While analyses of bulk tissues can reveal changes in chromatin accessibility, such changes often reflect shifts in cell composition rather than functional alterations within specific cell [19]. Moreover, bulk methods may fail to identify moderate but functionally important alterations in rare or specialized cell populations. To address this limitation, recent studies have also characterized cell-type-specific chromatin accessibility [20]. Single-cell chromatin accessibility analyses in the brain have provided deeper insight into epigenetic regulation at cellular resolution [21], including its role in AD pathogenesis [22]. For example, studies of neuronal and non-neuronal cells have identified regulatory genomic signatures associated with AD [23]. Resolving chromatin accessibility at the single-cell level has also uncovered cell identity loss that contributes to disease progression [24,25]. However, spatially resolved epigenetic analyses at the cellular level remain limited, particularly in capturing spatial epigenetic alterations in the brain of AD patients and mouse models.

Recently, spatial mapping of chromatin accessibility across tissue sections with the spatial assay for transposase-accessible chromatin using sequencing (spatial-ATAC-seq) technology has emerged as a powerful tool for delineating region-specific epigenetic landscapes and identifying gene regulators involved in central nervous system (CNS) development [26]. Additionally, spatially co-profiling chromatin accessibility and transcriptomes in mouse and human hippocampal tissue sections [27,28] has advanced the study of neurodegenerative diseases by offering a deeper understanding of the molecular basis of these conditions. Understanding the spatial distribution of epigenetic alterations in cell types across different brain regions holds significant potential for uncovering the complex molecular processes of human diseases, such as AD.

Chromatin accessibility is a critical determinant of gene regulation, influencing transcription factor binding and thereby shaping cell-specific responses to pathological stimuli such as Aβ. In this study, we employ spatial ATAC-seq to profile chromatin accessibility in the cortical and hippocampal sections of 5×FAD and C57BL/6J mice, enabling the characterization of both brain region-specific and cell-type-specific chromatin landscapes. We first define seven major brain cell types based on their spatially resolved open and active chromatin patterns, focusing specifically on four microglial and three astrocyte subtypes. We then spatially map the cell-type-specific chromatin landscape across key brain regions to reveal potential region-dependent shifts in cell composition and transcriptional regulatory activity in response to Aβ accumulation in 5×FAD mouse brain. Subsequently, we identify cell-type-specific enrichments in motifs and key transcription factors potentially involved in regulating gene expression changes. By integrating AD risk genes with our spatial ATAC-seq data, we pinpoint specific cell types exhibiting unique chromatin accessibility patterns related to these AD risk signals. Finally, we highlight microglia-specific chromatin accessibility at the *Vsir* gene locus as a potential therapeutic target for AD.

## Results

### Spatial profiling of chromatin accessibility in 5×FAD mouse brain

To gain comprehensive spatial insights into chromatin accessibility alterations in an AD mouse brain, we employed spatial ATAC-seq to characterize region- and cell-type-specific changes in the 6-month-old 5×FAD brain compared to C57BL/6J controls and link these changes to AD pathogenesis (**Fig. 1**). We analyzed sagittal brain sections from both 5×FAD AD model and C57BL/6J littermate mice, focusing on three major brain regions—the cortex, hippocampus, and thalamus—to capture region- and cell-type-specific chromatin accessibility changes. This analysis successfully sequenced 2,253 individual pixels (i.e., spatially barcoded regions) from the 5×FAD model and 2,345 pixels from the control (**Table S1**). Each tissue section achieved an average sequencing depth of 175.85 million reads, corresponding to approximately 76,488 reads per pixel. We obtained a median of 67,808 and 63,685 unique fragments per pixel in the 5×FAD and control samples, respectively. The average number of unique fragments detected and total mapped reads per pixel was comparable between the two samples (**Fig. S1A**). As expected, sequencing reads were predominantly mapped to transcription start sites (TSSs), with 3.54% and 3.32% of reads overlapping TSS regions, and 16.76% and 15.88% located in promoter peaks for the 5×FAD and control samples, respectively (**Fig. S1B**). Comparison of fragment size distributions and distances to transcription start sites (TSS) revealed high consistency between 5×FAD and control samples (**Fig. S1C, D**). The distribution of TSS enrichment scores versus unique nuclear fragments per pixel further confirmed the high quality of the data (**Fig. S1E**). These results show the high quality of the spatial ATAC-seq data, providing a foundation for downstream analysis of region- and cell-type-specific chromatin accessibility changes in the AD mouse brain.

**Fig. 1.**
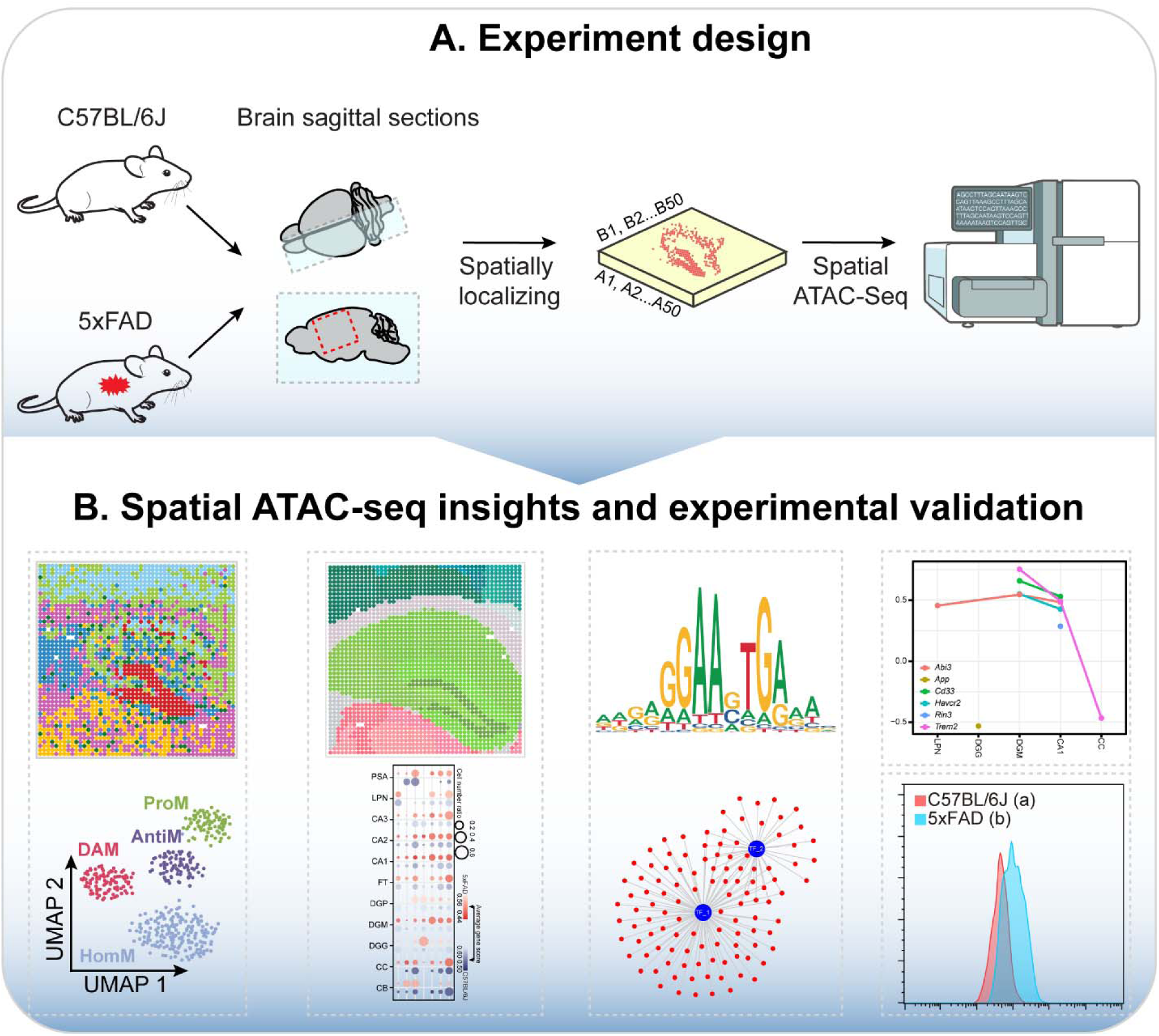
Experimental workflow and key findings from spatial ATAC-seq in 5×FAD and control mouse brains. (A) Schematic of the experimental design. Spatial ATAC-seq was performed on sagittal brain sections from 5×FAD and C57BL/6J mice to profile chromatin accessibility across defined brain regions. (B) Key findings from the study include: (1) identification of major brain cell types and glial subtypes; (2) region-specific chromatin accessibility patterns; (3) cell-type-specific transcription factor motif enrichment; (4) differential chromatin accessibility in AD risk genes; and (5) experimental validation of *Vsir* expression in microglial subtypes.

### Spatial chromatin accessibility mapping of the mouse brains

We next sought to identify brain region-specific and cell-type-specific changes in chromatin accessibility using inferred gene score, which quantifies chromatin accessibility across gene bodies and flanking regions (i.e., 5,000 bp upstream). The gene score incorporates two weighting factors: the distance of accessibility signals from the gene and the size of the gene itself (**see methods**). With inferred gene scores, we identified seven distinct clusters of spatial pixels, which were annotated using cell-type-specific gene markers (**Fig. 2A**). For simplicity and biological interpretability, we refer to spatial pixels as “cells” when discussing inferred cell-type-specific chromatin accessibility patterns, although the data are derived from pixel-level resolution. These included three neuronal subtypes (756, 1,002, and 385 pixels), oligodendrocytes (1,182 pixels), astrocytes (557 pixels), microglia (444 pixels), and granule cells (272 pixels) (**Fig. 2B**). These cell types were consistently present in both 5×FAD and control mice (**Fig. 2C; Fig. S2A**). Both glial cells, including microglia and astrocytes, were substantially increased in 5×FAD compared to controls, aligning with earlier findings in human AD samples [29] and 5×FAD mice [30,31] using single cell RNA sequencing (**Fig. S2B**).

**Fig. 2.**
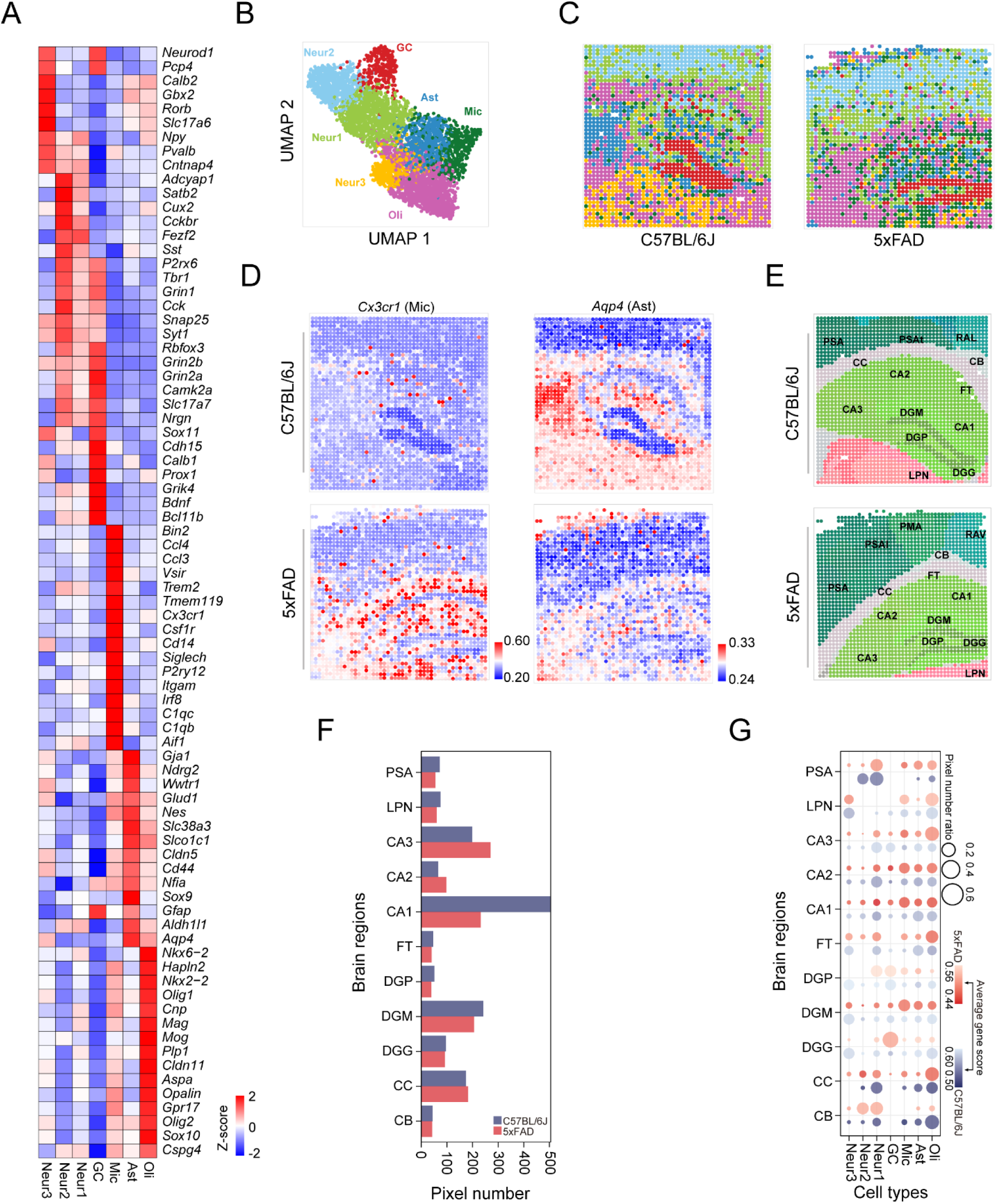
Spatial transcriptomic and cell-type-specific analysis in the brain of C57BL/6J and 5×FAD mouse model. (A) Heatmap of marker genes across various cell types, including neurons, astrocytes, oligodendrocytes, microglia, and granule cells. Rows represent genes, and columns correspond to cell types. (B) Uniform manifold approximation and projection (UMAP) plot showing clustering into distinct cell types, including three neuronal subgroups (i.e., Neuron1, Neuron2, Neuron3), astrocytes, microglia, oligodendrocytes, and granule cells. Each dot represents a single cell, colored by its corresponding cluster. (C) Spatial cell-type mapping of the hippocampus in C57BL/6J (left) and 5×FAD (right) mice. Each pixel area is color-coded according to its assigned cell type, demonstrating differences in cell-type composition and spatial organization between control and AD models. (D) Spatial expression patterns of selected marker genes across seven cell types: *Cx3cr1* (microglia), and *Aqp4* (astrocytes). (E) Anatomical registration of brain regions using the Allen Brain Atlas in C57BL/6J and 5×FAD mice. PSA (primary somatosensory area, barrel field, layer 6a), PMA (primary motor area, layer 5), RAV (retrosplenial area, ventral part, layer 5), PSAl (primary somatosensory area, lower limb, layer 5), RAL (retrosplenial area, lateral agranular part, layer 6a), PSAt (primary somatosensory area, trunk, layer 6a), CB (cingulum bundle), CC (corpus callosum), FT (fiber tracts), CA1 (field CA1), CA2 (field CA2), CA3 (field CA3), DGM (dentate gyrus, molecular layer), DGP (dentate gyrus, polymorph layer), LPN (lateral posterior nucleus of the thalamus), DGG (dentate gyrus, granule cell layer) (F) Bar plot comparing the number of pixels assigned to 11 annotated brain regions between C57BL/6J and 5×FAD mice. (G) Bubble plot showing the average gene scores across cell types within each brain region. Bubble size indicates pixel number; color intensity reflects average gene score for C57BL/6J (gray) and 5×FAD (red) mice.

As expected, specific brain cell types were mapped to distinct anatomical locations (**Fig. 2C**). Neuron 1 (Neur1, yellow green) are primarily located in the cortex and the CA1, CA2, and CA3 regions of the hippocampus. Neuron 2 (Neur2, pale blue) are found in CA2 and cortex, and neuron 3 (Neur3, amber) are mainly distributed throughout the thalamic region. Neuron 3 was markedly decreased in the thalamic region in 5×FAD compared to the control. Oligodendrocytes (Oli, vivid pink) are dispersed across white matter areas, supporting the presence of myelination in deeper cortical and subcortical regions. Astrocytes (Ast, steel blue) are primarily located around the hippocampus, with a small number distributed in the cortex, providing structural and metabolic support. Microglia (Mic, dark green) are scattered across most regions, with the majority concentrated around the hippocampus. Granule cells (GC, red) are specifically localized to the hippocampal dentate gyrus and are rarely observed in the CA1, CA2, and CA3 regions. These clusters exhibited unique chromatin accessibility patterns for cell-type specific marker genes (**Fig. 2D**), including neurons (*Fezf2*, *Slc17a6*), oligodendrocytes (*Olig1*), microglia (*Cx3cr1*), astrocytes (*Aqp4*), and granule cells (*Prox1*).

To generate a well-annotated spatial chromatin accessibility landscape and assess gene expression profiles across brain regions, we performed co-registration of spatial ATAC-seq cluster images to the Allen Mouse Brain Common Coordinate Framework (CCFv3) using the QUINT workflow. Given subtle structural differences in the brains used for spatial ATAC-seq profiling, images were registered separately (**Fig. 2E**). Regions of interest (ROIs; sub-regions) were defined based on anatomical references from the Allen Brain Atlas. We identified a total of 48 and 52 ROIs from 5×FAD and C57BL/6J control mice, respectively. Eleven brain regions containing more than 30 pixels were included for downstream analyses. The pixel numbers in each brain region were generally comparable between 5×FAD and control mice, with the exception of the hippocampal CA1 field (**Fig. 2F**). However, cell type composition and average gene activity scores varied substantially across brain regions (**Fig. 2G**). For instance, microglia were significantly increased in the CA1, CA2, and CA3 fields, as well as the dentate gyrus molecular layer between 5×FAD and control mice (**Fig. 2G**). Differential chromatin accessibility analysis between 5×FAD and control mice across these brain regions revealed subregion-specific genes (**Table S2**), such as *Oprk1* and *Neurl3* for CA1, *Nlrp3*, *Ccl4* and *Lyn* for CA2, *Ccl3* and *Cyp27a1* for CA3, *Ryr2* for dentate gyrus, granule cell layer (**Fig. S2B**). Enrichment analysis revealed significant activation of genes linked to immune response and cytokine signaling pathways in the CA1 and CA3 regions, but not in CA2 (**Table S3**). This pattern aligns with previous findings that CA1 is highly vulnerable to AD-related pathology, followed by CA3, while CA2 remains relatively resistant [32]. These results enabled robust mapping of the chromatin accessibility landscape across key brain regions, facilitating the identification of changes in regulatory element accessibility affecting gene expression.

### Chromatin accessibility landscape of microglia in the 5×FAD brain

Genetic analyses have revealed that many AD risk genes, such as *Trem2*, *CD33*, and *Bin1*, exhibit microglia-specific expression [33]. Furthermore, single-cell and single-nucleus sequencing studies have identified disease-associated microglia (DAM) in both human AD and 5×FAD mouse brains [34]. Using microglia marker genes (**Fig. 3A**), we classified microglia into four subtypes: homeostatic microglia (HomM), pro-inflammatory microglia (ProM), anti-inflammatory microglia (AntiM), and disease-associated microglia (DAM) (**Fig. 3B**). Our analysis revealed an increase in DAMs in 5×FAD brains, with predominant distribution in the hippocampal region (**Fig. 3C**). Differential gene activity score analysis identified 225 genes between 5×FAD and control samples, including 156 upregulated and 69 downregulated genes (**Fig. 3D, Table S4**). These genes include several known to be associated with AD, such as *Apoe*, *Lpl*, *Apoc1*, and *Col18a1* (**Fig. S3A, B**). These genes were enriched in pathways related to blood-brain barrier (BBB) integrity and cerebrovascular dysfunction, synaptic dysfunction and neuronal signaling, neuronal death and impaired neurogenesis, neuroinflammation and immune dysregulation, calcium dysregulation and neuronal homeostasis, oxidative stress and cellular stress response and were upregulated in AD brains (**Fig. 3E, Table S5**).

**Fig. 3.**
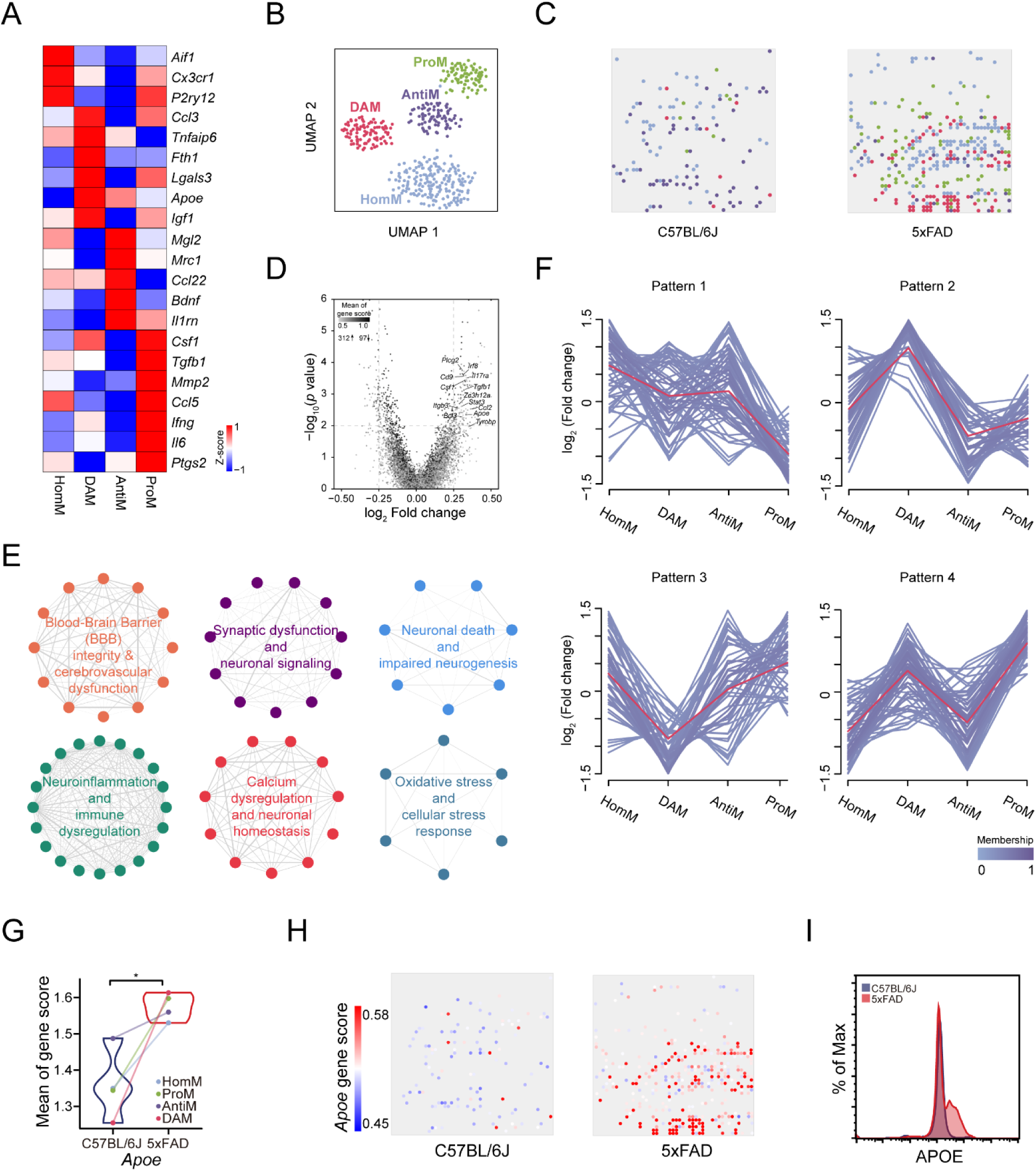
Characterization of microglial subtypes and their alterations in 5×FAD compared to control mouse brains. (A) Heatmap of marker gene expression across four microglial subtypes: homeostatic microglia (HomM), pro-inflammatory microglia (ProM), anti-inflammatory microglia (AntiM), and disease-associated microglia (DAM). Red indicates higher expression; blue indicates lower expression relative to other subtypes. (B) UMAP plot showing distinct clustering of the four microglial subtypes based on gene expression profiles. (C) Spatial distribution of microglial subtypes in sagittal brain sections from C57BL/6J (left) and 5×FAD (right) mice, revealing changes in subtype composition and localization. (D) Volcano plot displaying differentially expressed genes (DEGs) between 5×FAD and control microglia. Red and blue dots represent significantly upregulated and downregulated genes, respectively; gray indicates non-significant genes. (E) Gene ontology (GO) enrichment analysis highlighting six major functional clusters associated with DEGs, including neuroinflammation, synaptic dysfunction, and oxidative stress. (F) Mfuzz clustering of gene score patterns across the four microglial subtypes, revealing four distinct gene expression trajectories. (G) Comparison of *Apoe* gene scores across microglial subtypes in C57BL/6J and 5×FAD mice; asterisks indicate significant differences. (H) Spatial expression maps of *Apoe* in C57BL/6J (left) and 5×FAD (right) brains. Color scale represents gene score intensity. (I) Experimental validation showing increased APOE expression in 5×FAD microglia compared to controls, based on flow cytometry or sequencing data.

The dynamics of microglial subtypes by the pseudotime analysis revealed a transition of microglia from a HomM state to DAMs, AntiM, and ProM (**Fig. S3C**). To further explore the molecular changes associated with this transition, we examined distinct gene expression patterns across microglial subtypes (**Fig. 3F, Table S6**). Pattern 1 shows relatively stable expression across HomM, DAM, and AntiM, but a pronounced reduction in ProM. This pattern may reflect genes that are downregulated during the transition to a pro-inflammatory phenotype and could play roles in maintaining basal microglial function. Representative genes in this pattern include *Selplg*, *Ctsd*, and *Irf8*. Pattern 2 is characterized by a peak in DAM, with reduced expression in the other states, suggesting involvement in early microglial activation. This further suggests their potential role in early microglial activation and neuroinflammatory responses, including altered expression of genes such as *CD68*, *Apoe* and *Gps2*. Pattern 3 genes exhibit a sharp downregulation in DAM, followed by a rebound in AntiM and ProM, forming a U-shaped trajectory. This indicates transient suppression during the disease-associated state, with reactivation in later phases of microglial polarization. These genes may contribute to cellular recovery or adaptive remodeling, including *Mdp1*, *Vsir*, and *Rgcc*. Pattern 4 showed upregulation in both DAM and ProM, suggesting involvement in early and sustained phases of microglial activation—potentially through lipid remodeling or pro-inflammatory signaling pathways—with representative genes including *Atat1*, *Lcat*, and *Kctd11*.

We found that chromatin accessibility at the *Apoe* locus—an established AD risk gene— was elevated in 5×FAD compared to the C57BL/6J control samples across all four microglial subtypes (HomM, ProM, AntM, and DAM) (**Fig. 3G**). Spatial maps further highlighted upregulation of APOE chromatin accessibility in the 5×FAD brain, primarily in hippocampal regions and to a lesser extent in cortical areas (**Fig. 3H**). Complementary flow cytometry analysis confirmed increased APOE protein expression in microglia from 5×FAD mice, evidenced by a rightward shift in the fluorescence intensity distribution relative to controls (**Fig. 3I**). These results indicate that APOE is transcriptionally upregulated across four microglial subpopulations in the AD mouse brain, suggesting a potential role in driving disease-associated microglial reprogramming.

### Chromatin accessibility of Astrocytes in the 5×FAD brain

Astrocytes, another type of glial cells in the brain, play essential roles in maintaining neural homeostasis, supporting neuronal function [35]. Recent studies have shown that the alteration of astrocytes is involved in the initiation and progression of AD [36,37]. To further investigate astrocyte function in the spatial context of the AD brain, we analyzed chromatin accessibility in astrocytes. Using enrichment expression of marker genes (**Fig. 4A**), we classified astrocytes into three subtypes: homeostatic astrocytes (HomA), proinflammatory and neurotoxic A1-reactive astrocytes (A1rA), and A2-like reactive astrocytes (A2rA) (**Fig. 4B**). Unsupervised clustering revealed distinct separation of these astrocyte subtypes, with homeostatic astrocytes predominantly found in control brains, whereas A1 and A2 reactive astrocytes were more abundant in 5×FAD brains (**Fig. 4C**).

**Fig. 4.**
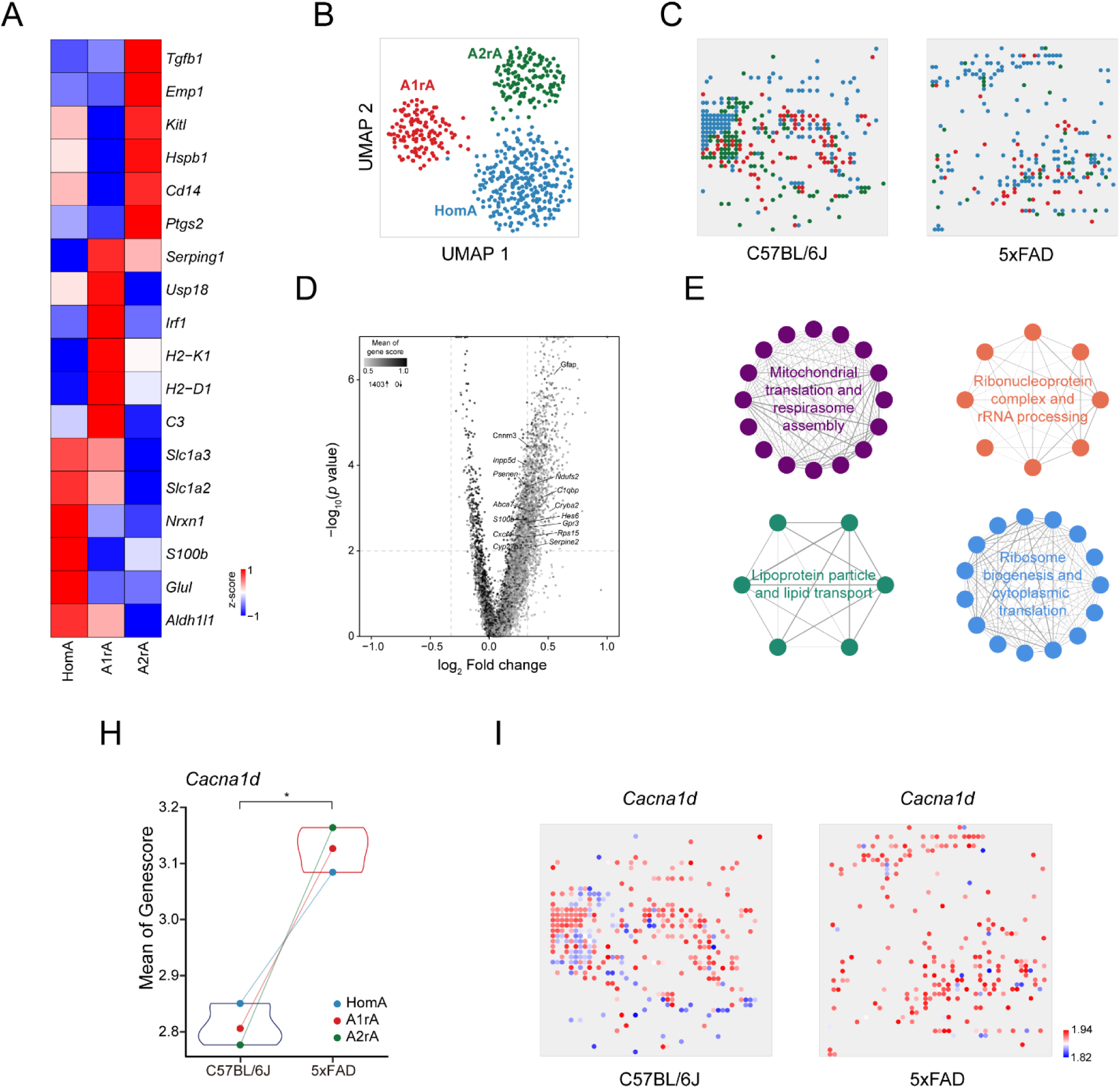
Analysis of astrocyte subtypes and alteration in 5×FAD compared to control mouse brains. (A) Heatmap displaying marker gene expression across three astrocyte subtypes: homeostatic astrocytes (HomA), A1 reactive astrocytes (A1rA), and A2 reactive astrocytes (A2rA). Red indicates higher relative expression, while blue indicates lower expression. (B) UMAP plot showing clustering of astrocyte subtypes based on gene expression profiles. (C) Spatial distribution of astrocyte subtypes in sagittal brain sections of C57BL/6J (left) and 5×FAD (right) mice, revealing shifts in subtype composition and localization. (D) Volcano plot illustrating genes with differential chromatin accessibility between C57BL/6J and 5×FAD astrocytes. Red-labeled genes represent significant upregulation in 5×FAD. (E) Functional enrichment analysis of differentially accessible genes, highlighting pathways related to lysosomal transport, mitochondrial dysfunction, synaptic signaling, and proteasomal degradation. (H) Gene score comparison for *Cacna1d* across astrocyte subtypes in 5×FAD and control mice. Asterisks denote statistically significant differences. (I) Spatial expression maps of *Cacna1d* in C57BL/6J (left) and 5×FAD (right) brain sections. Color scale represents gene score intensity.

Differential chromatin accessibility analysis identified several genes with altered accessibility between 5×FAD and control astrocytes (**Fig. 4D, Table S7**), including *Csf1r*, *Tspo*, and *Vwf* (**Fig. S4**) Functional enrichment of these genes highlighted pathways related to synaptic dysfunction, lysosomal and proteasomal pathways, lipoprotein metabolism, and mitochondrial dysfunction, all of which are implicated in AD pathology (**Fig. 4E, Table S8**). For example, *Cacna1d*, a gene involved in calcium channel function, was found to have significantly increased accessibility in 5×FAD astrocytes compared to controls (**Fig. 4H**). Spatial mapping of *Cacna1d* chromatin accessibility showed distinct regional differences, with increased accessibility in high-vulnerability, disease-associated regions of 5×FAD brains compared to controls (**Fig. 4I**).

### Cell-type-specific alterations of transcription factors in the brain of 5×FAD

Many transcription factors (TFs), such as ATF4, have been implicated in AD [38]. To explore TFs in a cell-type-specific context, we examined the enrichment for transcription-factor-binding motifs in altered chromatin accessible regions across major brain cell types. Our analysis revealed distinct patterns of TF enrichment (**Fig. 5A**). In microglia, binding motifs for TFs such as *Sfpi1, Bcl11a, Bcl11b, Elf3,* and *Elf5* were enriched in areas of open chromatin. For example, the top enriched TF, *Sfpi1* (encoding PU.1) (**Fig. 5B**), is a known AD risk gene [39,40], regulating downstream expression of microglial genes involved in phagocytosis and immune responses [41]. Prior studies have shown that *Spi1* knockdown exacerbates AD-related pathology, including Aβ aggregation, plaque accumulation, and gliosis, while *Spi1* overexpression confers protection against these phenotypes [42]. Astrocytes exhibited enrichment for TFs such as *Nfix, Egr1, Zfp263,* and *Wt1*. Among these, Egr1 has been shown to be regulated by APP via acetylation of histone H4 at its promoter region [43]. Furthermore, *Egr1* promotes Aβ production by transcriptionally activating *BACE-1* [44] and *PSEN2* [45], both of which are key AD risk genes.

**Fig. 5.**
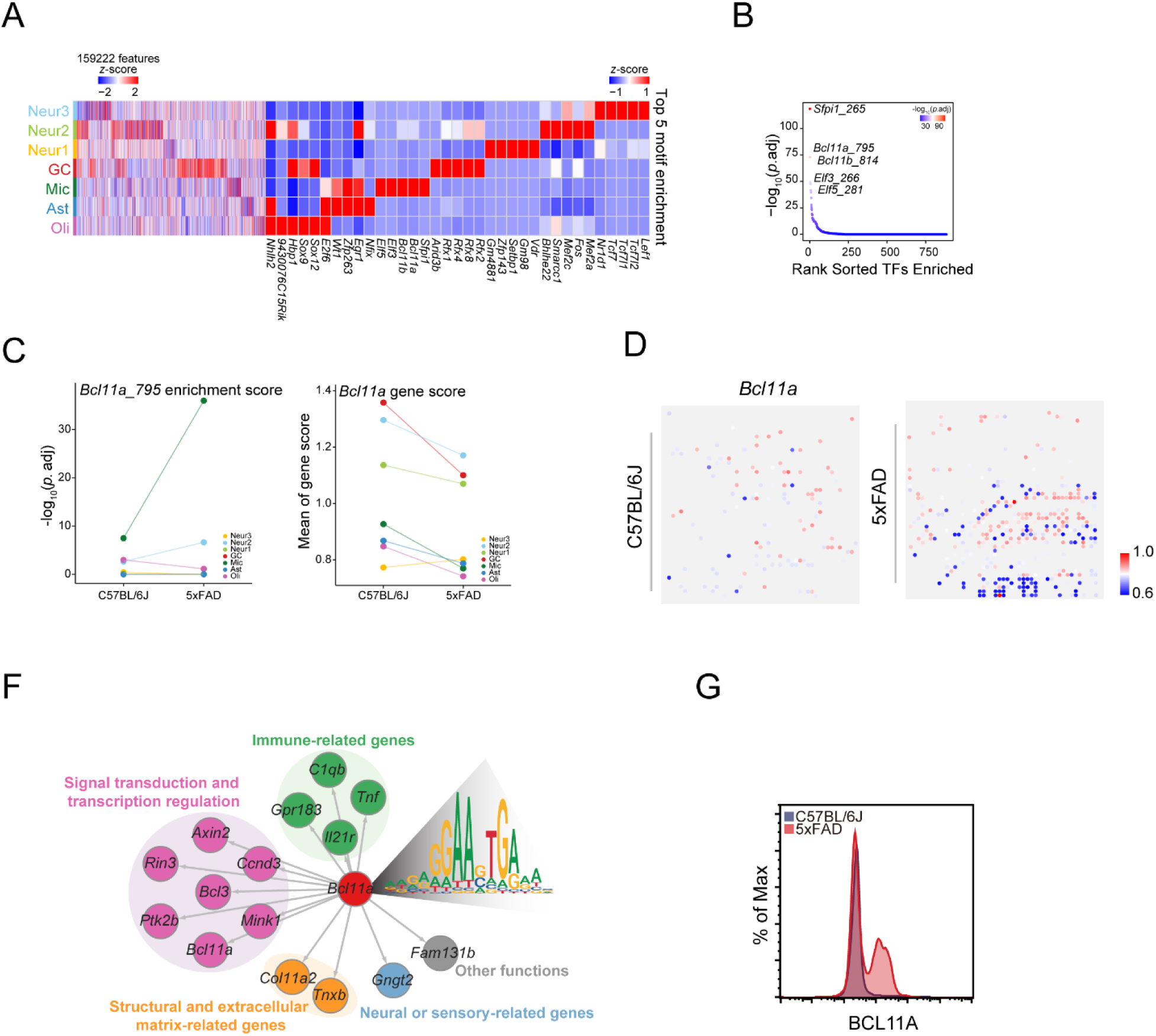
Cell-type-specific transcription factor activity and *Bcl11a* regulation in microglia. (A) Heatmap showing transcription factor (TF) motif enrichment and chromatin accessibility across 159,222 ATAC-seq peaks. The left panel displays peak accessibility (Z-scores) across major brain cell types—granule cells (GC), neurons (Neur1–3), microglia (Mic), astrocytes (Ast), and oligodendrocytes (Oli). The right panel highlights the top five enriched TF motifs for each cell type. (B) Ranked plot of transcription factor enrichment scores in microglia, with *Bcl11a* among the top-enriched TFs. (C) Comparisons of *Bcl11a*-related enrichment scores (top) and gene scores (bottom) in C57BL/6J versus 5×FAD microglial populations. (D) Spatial gene score maps of *Bcl11a* in C57BL/6J (top) and 5×FAD (bottom) mouse brains, showing cell-type- and region-specific expression patterns. (E) Spatially localized expression of *Bcl11a* at the single-cell level in C57BL/6J (left) and 5×FAD (right) mice. (F) Predicted *Bcl11a* regulatory network. Downstream target genes are grouped by functional categories, including immune-related, neural/sensory-related, transcriptional regulators, and extracellular matrix-related genes. The *Bcl11a* motif is also shown to the right. (G) Flow cytometry validation showing increased expression of *Bcl11a* in microglia from 5×FAD mice compared to controls.

Additionally, *Bcl11a* emerged as the second enriched regulator in microglia (**Fig. 5B, C**). It exhibits a significant elevation in motif enrichment in 5×FAD compared to control, but not for the gene score (**Figure 5C, D**). Regulatory network analysis showed that *Bcl11a* regulates 15 downstream target genes with promoter-associated accessibility peaks, including *Rin3* and *Ptk2b* (**Fig. S5**), which are involved in immune-related pathways, signaling pathways, transcription regulation, and structural/extracellular matrix-related genes (**Fig. 5E**). Although direct links between *Bcl11a* and AD are limited, *Bcl11a* has been implicated in several neurological disorders, including Huntington’s disease, autism spectrum disorder, and schizophrenia (41). Flow cytometry analysis further confirmed that BCL11A expression was notably increased in microglia from 5×FAD mice compared to those from control mice (**Fig. 5F**). These findings highlight BCL11A as a key transcriptional regulator in AD.

### Alterations in chromatin accessibility associated with AD risk genes

While it is well-established that genetic risk variants for AD are enriched in noncoding regions and regions of open chromatin [46], their quantitative relationship with chromatin accessibility across specific brain regions and cell types remains unclear. To address this, we focused on 38 AD-associated genes identified in a recent GWAS [47]. Among these, 1,094 genes exhibited significant differences in gene scores, with six AD risk genes—*Trem2*, *Abi3*, *App*, *Cd33*, *Havcr2* and *Rin3* — showing notably altered chromatin accessibility (**Fig. S6**). These six AD risk genes were significantly overrepresented among the differentially accessible genes, (*p*-value = 7 x 10^-3^; Fisher’s exact test). *Trem2* displayed a region-specific pattern, with elevated gene scores in the hippocampus and a marked reduction along the granule cell (GC) to cortex axis (**Fig. 6A**). As anticipated, *Trem2* accessibility was significantly increased in microglia, with no observable changes in other cell types (**Fig. 6B**). Spatial mapping showed a similar pattern as microglial populations and revealed enhanced *Trem2* accessibility in 5×FAD, particularly in hippocampal regions, compared to controls (**Fig. 6C, D**). Chromatin accessibility tracks revealed an increase in an open chromatin peak near the TSS of the *Trem2* locus in microglia from 5×FAD mice (**Fig. 6E**). Motif enrichment and transcription factor network analysis suggested *Stat1* as a potential upstream regulator of *Trem2*, along with other associated regulators, such as *Nr4a1, Nr4a2, Nr4a3, Stat6,* and *Mocp3*, all enriched in microglial populations (**Fig. 6F**). Taken together, this analysis highlights region- and cell type-specific epigenetic alterations in AD risk genes, with *Trem2* emerging as a key microglial target of chromatin remodeling in AD.

**Figure 6.**
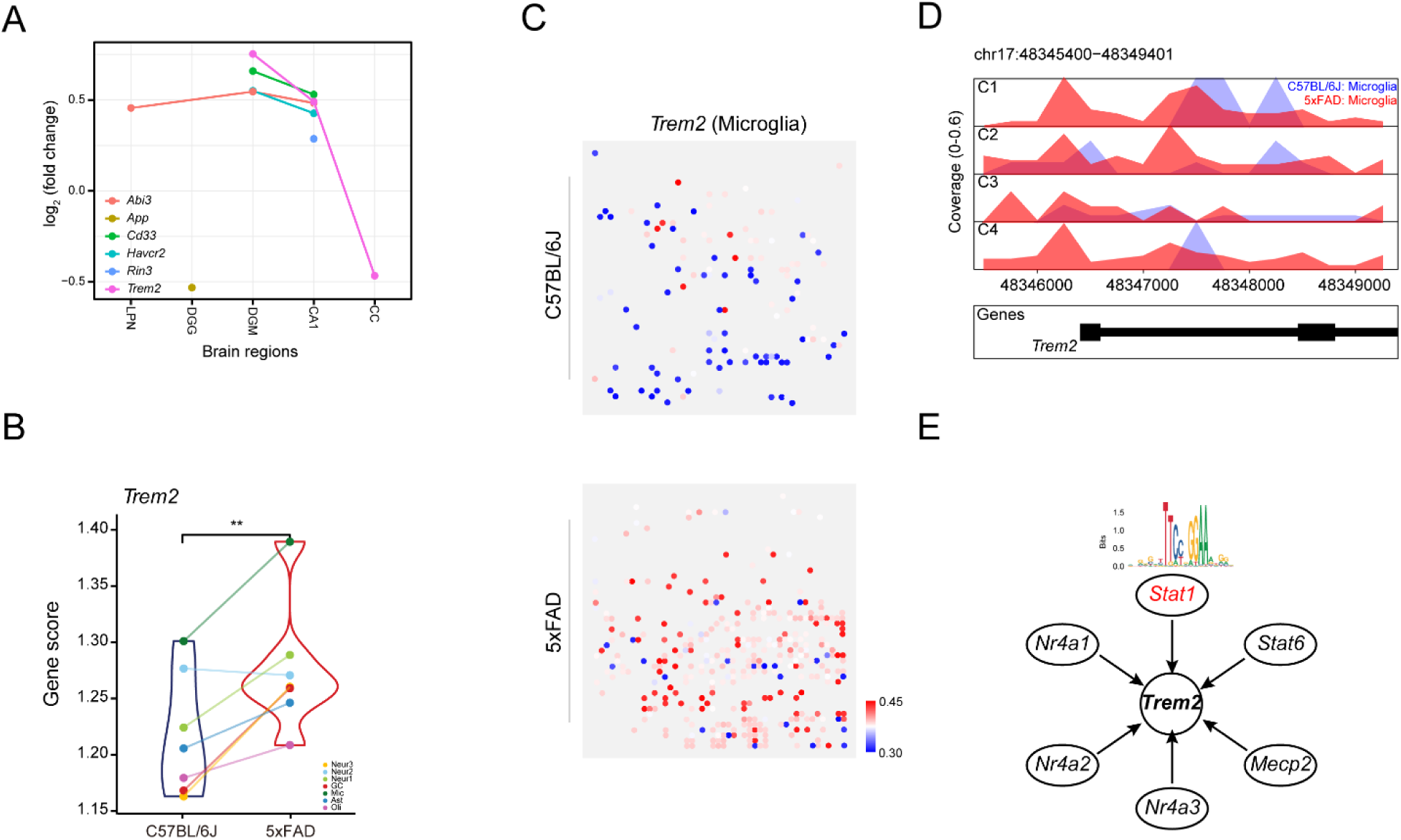
Alterations in chromatin accessibility associated with *Trem2*, an AD risk gene. (A) Trend of gene scores for 6 AD risk genes across major brain regions from inside to outside (LPN, DGG, DGM, CA1, and CC). *Trem2* shows a significant decrease in CC compared to other regions. (B) Violin plot showing significantly increased *Trem2* gene score in microglia in 5×FAD compared to C57BL/6J. (C) Cell-type–specific spatial mapping of *Trem2* gene scores within microglial populations. Elevated Trem2 signal is observed in 5×FAD brains compared to controls. (D) Chromatin accessibility signal tracks for *Trem2* locus across cell types, showing enhanced peaks in microglia from 5×FAD brains. Genome coordinates based on mm10. (E) Transcriptional regulatory network centered on *Trem2*, inferred from motif enrichment analysis. *Stat1* appears as a key upstream regulator with increased motif activity in 5×FAD microglia.

### VISTA dysregulation across microglial subtypes in 5×FAD

Neuroinflammation driven by microglial activation is one of the hallmarks of AD [48,49]. Dysregulation of immune checkpoint pathways may contribute to disease pathogenesis and progression by altering microglial activation [50]. Negative checkpoint receptors (NCRs) play a critical role in maintaining microglial homeostasis and regulating immune responses [50]. Among these NCRs, a particularly notable one is the V-domain Ig suppressor of T cell activation (VISTA). The chromatin regions of the VISTA locus were more accessible in microglia than in other cell types (**Fig. 7A**). Microglia from 5×FAD exhibited notably increased chromatin accessibility at the VISTA locus compared to controls (**Fig. 7A**). Moreover, a greater number of brain regions displayed increased chromatin accessibility at the VISTA locus in 5×FAD mice (**Fig. 7B**). Microglia-specific analysis revealed that chromatin accessibility in the promoter region at *Vsir* locus is elevated in 5×FAD (**Fig. 7C**).

**Fig. 7.**
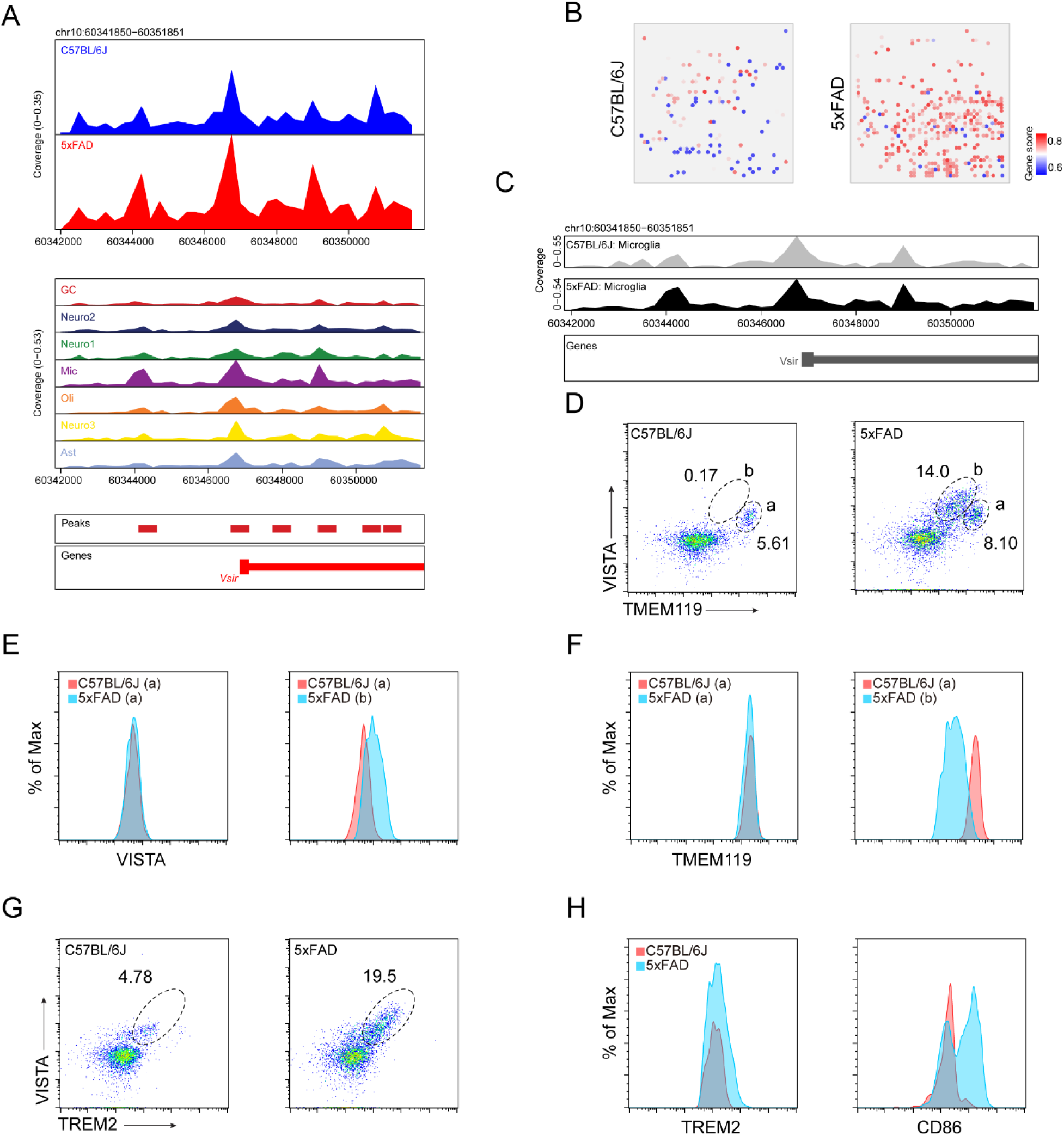
Chromatin accessibility of the Vsir (VISTA) gene in control and 5×FAD mouse brains. (A) Genome browser tracks showing chromatin accessibility at the *Vsir* locus in the Psudo-bulk level (top), and across seven cell types (middle). A total of six peaks within the region of *Vsir* locus were used for calculating gene score (bottom). (B) Chromatin accessibility of *Vsir* locus specifically in microglia from C57BL/6J (gray) and 5×FAD (black) mice. (C) Gene scores of *Vsir* in each microglial state for C57BL/6J (gray) and 5×FAD (black) mice. (D) Flow cytometry plots showing increased percentages of VISTA⁺TMEM119⁺ and VISTA⁺TMEM119⁻ microglia in 5×FAD mice compared to C57BL/6J controls. (E) Histogram plots showing increased *Vsir* protein (VISTA) expression in microglia from 5×FAD mice across two biological replicates. (F) Flow cytometry analysis of TMEM119 expression, which remains relatively unchanged between 5×FAD and control mice. (G) Co-staining of VISTA and TREM2 reveals a substantial increase in VISTA ⁺TREM2⁺ microglia in 5×FAD mice. (H) Flow cytometry histograms showing elevated TREM2 and CD86 protein expression in 5×FAD microglia, supporting a DAM-like, pro-inflammatory phenotype.

To validate *Vsir* expression changes observed in our spatial ATAC-seq data, we performed flow cytometry analysis on microglia isolated from C57BL/6J and 5×FAD mice. Consistent with chromatin accessibility findings, VISTA protein expression was markedly increased in microglia from 5×FAD mice compared to controls (**Fig. 7D–E**). Co-expression analysis showed a greater proportion of VISTA^+^TMEM119^+^ microglia in 5×FAD mice (14.0%) relative to C57BL/6J mice (0.17%), with an additional shift in a distinct VISTA^+^TMEM119^-^ population (8.10% vs. 5.61%). In contrast, TMEM119 expression remained relatively unchanged (**Fig. 7F**), supporting a selective increase in VISTA. Co-staining with TREM2 further revealed enrichment of the VSIR^+^TREM2^+^ subset in 5×FAD microglia (19.5% vs. 4.78%; **Figure 7G**), suggesting a potential link between *Vsir* upregulation and DAM-like phenotypes. Additional profiling showed increased expression of other activation markers, including CD86 (**Fig. 7H**), further indicating a shift toward a pro-inflammatory state.

## Discussion

In this study, we employed spatial-ATAC-seq to generate spatially resolved profiles of chromatin accessibility in the brains of an AD mouse model. To our knowledge, this is the first study to spatially profile and analyze chromatin accessibility in the 5×FAD mouse brain. We identified region- and cell-type-specific patterns of chromatin accessibility and characterized their alterations between 5×FAD and control mice. Pathway enrichment analysis of cell-type-specific differentially accessible regions revealed activation of immune response and neuronal signaling pathways in microglia, and secretion- and neurotransmission-related pathways in astrocytes. Moreover, we identified cell-type-specific gene regulators and linked changes in chromatin accessibility to AD risk genes, providing a detailed epigenetic landscape of AD. We finally validated increased expression of *Vsir* in microglia using flow cytometry. Collectively, these findings underscore the importance of chromatin accessibility in deciphering the molecular mechanisms and changing landscape of gene expression that drives AD pathogenesis.

The architecture of brain regions is intricately linked to their function, and disruptions in this spatial organization have been implicated in AD [51]. For example, neurons in layers II and III of the cortical regions, as well as those in the hippocampal formation, are particularly vulnerable to AD pathology due to the early accumulation of Aβ [52]. The interactions between neurons, microglia, and astrocytes are critical for maintaining homeostasis, and growing evidence suggests that dysregulated neuron–glia communication contributes to synaptic dysfunction, neuroinflammation, and progressive neurodegeneration in AD [53]. Spatial omics technologies offer an opportunity to uncover cell-type-specific changes within the critical brain regions. In this study, we found that the hippocampal CA1 region exhibited the greatest number of genes with altered chromatin accessibility between 5×FAD and control mice (**Table S2**). Interestingly, the cohort of accessible genes in the CA1 were enriched predominantly for regulation of cytokine production and canonical NFκβ signal transduction (**Table S3**).

Glial cells are increasingly recognized as key players in the pathogenesis of AD as their roles in neuroinflammation, synaptic pruning, and metabolic homeostasis [54]. Despite the importance of glial cells and crosstalk with neuronal cells, fully understanding the functional roles of the glial cells remains a major challenge, largely due to the complex cellular heterogeneity within the brain and the dynamic nature of disease progression. Recent advances in single-cell and single-nucleus RNA sequencing have begun to unravel this complexity, revealing distinct microglial and astrocyte subpopulations associated with disease states in both mouse models and human brain tissues [55,56]. However, the spatial distribution and functional relevance of these subpopulations across different brain regions remain poorly understood. In this study, we found that the DAM subpopulation is more abundant in hippocampal tissues (**Fig. 3C**), a key site of vulnerability in AD. The chromatin accessibility of DAM-related genes, such as *Apoe*, is also significantly elevated (**Fig. 3G–I**), suggesting enhanced activation of microglia in response to local pathological signals (e.g., Aβ stimuli). These findings support the notion that regional microglial responses may shape disease progression in a spatially resolved manner.

The epigenetic mechanisms—particularly chromatin accessibility—that govern their cell-type-specific transcriptional programs in the context of AD remain poorly understood. Understanding how chromatin landscapes differ across glial subtypes, especially microglia and astrocytes, can shed light on the molecular transitions from homeostatic to reactive states. Our spatial-ATAC-seq analysis reveals distinct patterns of chromatin remodeling in glial populations, suggesting that epigenomic reprogramming underlies their functional heterogeneity and disease-associated phenotypes. For example, the key AD risk gene *Trem2* exhibited a gradual decrease in chromatin accessibility (gene score) from the DG to CA1 and further to the CC regions (**Fig. 6A**). Therefore, defining regulatory changes is essential for identifying key transcriptional drivers of glial dysfunction and could lead to the identification of novel therapeutic molecules targeting glial-mediated processes in AD.

Like *Trem2*, *Vsir* functions as a cell surface receptor on microglia, promoting microglial chemotaxis and phagocytosis [57]. As a negative checkpoint regulator (NCR), *Vsir* plays a crucial role in regulating immune responses in the brain by inhibiting T-cell activation [58]. VISTA encoded by *Vsir* was shown to be enriched in TREM2 microglial subpopulation in 5×FAD, suggesting a potential link between *Vsir* upregulation and DAM-like phenotypes. These findings suggest that VISTA plays a complex, subtype-specific role in microglial-T cell interactions, influencing the neuroimmune landscape in AD. Further investigation into the mechanisms underlying these changes may offer new therapeutic strategies aimed at modulating VISTA activity to mitigate excessive neuroinflammation and its detrimental effects on disease progression.

A key limitation of our spatial ATAC-seq profiling is its inability to achieve single-cell resolution. In this study, we used a 50 μm pixel array, which captures signals from approximately 10–25 cells in the brain [59], making it difficult to distinguish different cell types within individual pixels. As a result, we defined only seven major cell types, primarily reflecting brain-region-specific chromatin accessibility. To overcome this limitation, future studies can integrate single-cell ATAC-seq with spatial ATAC-seq to computationally deconvolute mixed-cell signals [60,61]. Additionally, emerging high-resolution spatial profiling techniques, such as Visium HD and Stereo-seq, offer the potential to achieve single-cell resolution [62]. Computational approaches, including deep learning models and cell-type deconvolution algorithms, may further refine cell-type specificity in spatial datasets, enhancing our ability to resolve chromatin accessibility at the single-cell level.

In conclusion, we comprehensively characterized brain region-specific, cell-type-specific, and sub-cell-type-specific alterations in chromatin accessibility in the 5×FAD mouse model, highlighted cell-type-specific enrichment of transcription factors, and associated chromatin accessibility with AD risk genes. Our findings underscore the power of spatially resolved epigenomic approaches in unraveling the mechanisms of AD pathogenesis and provide an avenue to identify potential therapeutic targets, such as *Vsir*. Future studies that integrate additional layers of spatial multi-omic modalities, including transcriptomics and proteomics, will be essential for capturing cell-specific changes in the brain regions and further elucidating disease mechanisms.

## Methods and Materials

### Mouse brain samples

All animal experiments were approved by the Institutional Animal Care and Use Committee at University of North Dakota and performed in accordance with institutional guidelines. 5×FAD and C57BL/6J were obtained from the Jackson Laboratory. Six-month-old female mice were used in the experiments. All mice were maintained in a controlled environment of 22-23 °C, 50-60% humidity, 12□h alternating light-dark cycles, and with food and water provided *ad libitum*.

### Preparation of tissue slides

The flash frozen whole brain was dissected and sectioned at a thickness of 10□μm using a cryostat (Leica 3050S). Each of the serial sagittal sections of a brain region were placed onto an ultraclean glass slide (Electron Microscopy Sciences, 63478-AS). We took one slide for sequencing.

### Spatial-ATAC-seq profiling, barcoding, library preparation, and sequencing

Spatial ATAC-seq (AXO-0303) was performed following AtlasXomics’ protocols [63]. Briefly, tissue sections were fixed with 0.2% paraformaldehyde (PFA) for 5 minutes, quenched with glycine, and air-dried. Permeabilization was carried out using 0.2% NP-40 for 15 minutes, followed by a 30-minute tagmentation step under optimized conditions [63]. Tagmentation was then halted, and the tissue was washed and dried in preparation for spatial barcoding. ATAC-seq libraries were prepared using standard NGS amplification protocols. Single-index, 150□×□150 bp paired-end sequencing was performed on a NovaSeq 6000 (Illumina, 20012850) or NextSeq 2000 (Illumina, 20038897) with 15% PhiX spike-in for normalization across samples.

### Data preprocessing

To filter Read 2, two constant linker sequences (Linker 1 and Linker 2) were used. Integrated genome sequences were assigned to new Read 1, while barcodes A and B were incorporated into new Read 2. The resulting FASTQ files were aligned to the mouse reference genome (mm10), deduplicated, and quantified using Cell Ranger ATAC v2.0.0. BED-like fragment files were generated for downstream analysis. These fragment files contain information on genomic coordinates and spatial tissue locations, represented by the combination of barcode A and barcode B. We converted the input ATAC-seq fragment files into Arrow files using the “createArrowFiles” function in the ArchR program [64], with a minimum of 1,000 fragments per cell, a TSS enrichment filtering threshold of 4, and with the gene score matrix computation function enabled. The resulting Arrow files were used to initialize an ArchRProject with the appropriate genome annotation for subsequent analyses.

### Gene score calculation by ArchR

Gene score, defined as inference of gene expression from chromatin accessibility, was calculated using the ArchR package with a custom distance-weighted accessibility model. A default tile size of 500 bp was used to tile the genome, and tiles overlapping user-defined gene windows (default 100 kb) were considered. In this study, the promoter and gene body regions were included in the analysis. For each gene, ArchR selected tiles within the defined window that did not overlap with neighboring gene regions. The distance of each tile from the transcription start site was converted into a distance weight using the default model: e^(-|distance|/5000)^ + e^-1^

To account for differences in gene length, ArchR was used to apply an inverse gene size weight (1 / gene size), scaled linearly from 1 to 5. The analysis generated a gene score matrix for downstream analysis.

### Dimensional reduction, clustering, and marker identification

Dimensionality reduction was performed using iterative Latent Semantic Indexing (LSI) through the “addIterativeLSI” function, specifying 25,000 variable features, 25,000 final sampled cells, and two iterations. UMAP embedding was generated using the “addUMAP” function with minimum distance set to 1 for 40 defined nearest neighbors. Clustering was performed with “addClusters” at a resolution of 0.4 to resolve distinct cell populations. To enhance visualization and interpretation of gene activity patterns, MAGIC-based imputation was applied using “addImputationWeights”, allowing imputed features to be projected onto the UMAP. Marker genes and regions were identified using the “getMarkerFeatures” function with the “GeneScoreMatrix”, while correcting for technical biases including TSS enrichment and the log-transformed number of fragments. The Wilcoxon rank-sum test was used to assess statistical significance. Cell type identities were manually assigned by comparing canonical marker genes to cluster-specific markers.

### Registration of spatial ATAC-seq imaging data with the Allen Brain Atlas

To assign spatial ATAC-seq data to anatomical brain regions, we registered spatial ATAC-seq cluster images to the Allen Mouse Brain Common Coordinate Framework (CCFv3) [65]. The whole-brain image series were aligned to the Allen Brain Atlas using QuickNII [66], generating customized atlas maps that matched the cutting plane and proportions of each section. Barcodes from spatial ATAC-seq data were then extracted and assigned to individual brain regions using an in-house script.

### Pseudotemporal trajectory analysis for microglial subtypes

Pseudotemporal reconstruction was implemented by trajectory analysis using ArchR. A trajectory backbone was first created in the form of an ordered vector of cell group labels. We then used the “addTrajectory function” to create a trajectory and added the pseudo time to the UMAP projection.

### Analysis of differential chromatin accessible genes (DAGs)

To identify differentially accessible genomic regions, we performed the analysis of DAGs based on gene activity scores derived from chromatin accessibility data. Specifically, the GeneScoreMatrix, which quantifies chromatin accessibility across gene bodies and upstream regulatory regions, was extracted at the spatial pixel level and stored as a matrix representing inferred transcriptional activity across the tissue. Pairwise comparisons between 5×FAD and control samples were conducted using the Wilcoxon rank-sum test. We focused the analysis on protein-coding genes whose mean gene score across all pixels exceeded the median gene score of the entire dataset. Genes with a *p*-value < 0.01 and an absolute log₂ fold change greater than two times the global standard deviation were considered significantly differentially accessible. Patterns of differentially accessible genes (DAGs) in microglia were conducted using the Mfuzz [67] R package. We used the gene score of each cluster of 5×FAD samples minus that of the control samples.

### Enrichment analysis of DAGs

Enrichment analysis of DAGs was performed using two complementary tools: WebGestalt [68] and Metascape [69]. WebGestalt (http://bioinfo.vanderbilt.edu/webgestalt/) was used to perform Kyoto Encyclopedia of Genes and Genomes (KEGG) pathway enrichment analysis. Pathways with a *p*-value < 0.05 were considered statistically significant. Metascape (www.metascape.org) was used to cluster enriched terms and visualize functional groupings. The analysis was performed using the following parameters: minimum protein overlap of 3, enrichment *p*-value < 0.05, and at least 3 proteins per enriched group. Pathway visualizations were generated using Cytoscape version 3.10.2 [70].

### Motif enrichment analysis

Motif enrichment analysis was performed to identify TF binding motifs associated with differentially accessible chromatin regions. Motif enrichment for each cell type was then conducted using the “peakAnnoEnrichment” function with position weight matrices (PWMs) from the cisbp database provided in ArchR. Enrichment was calculated by comparing the frequency of motifs in the input peak set to a background set matched for GC content and peak length. Transcription factor motifs with adjusted *p*-values < 0.05 were considered significantly enriched in chromatin accessible regions. Results were visualized as enrichment heatmaps for each cell type.

### Identification of cell-type-specific enriched TFs

To identify which of these correlated TF regulators had strong differential motif activity differences, we calculated the average motif deviation scores with “exportGroupSE” in ArchR for each cluster and computed the max observed deviation difference between any two clusters. This motif difference and the TF-to-gene score correlation was then used to identify positive regulators (correlation > 0.5 and a maximum deviation score difference > 50th percentile). To construct the regulatory network between *Bcl11a* and its downstream genes, we integrated data from motif-peak and peak-to-nearest-gene annotations to identify transcription factor-gene associations, providing insights into their regulatory relationships with the data visualized using Cytoscape [70].

### Integration of cell-type-specific AD risk genes with DAGs

We integrated spatial cell-type-specific chromatin accessibility with the largest AD GWAS study published [7]. A total of 38 AD risk candidate genes were intersected with 1,094 DAGs identified between 5×FAD and control samples. The AD risk genes with differential chromatin accessibility were intersected with 6 TFs to identify AD-associated TFs potentially regulating disease-related gene networks.

### Isolation of microglia from rodent brains

C57BL/6J Wild-type (WT) and 5×FAD mice were euthanized using CO₂ and perfused with ice-cold PBS. Brains were rapidly collected and placed in a glass homogenizer containing 5 mL of dounce buffer (15 mM HEPES and 0.5% glucose in HBSS) supplemented with 500 units/mL DNase and 10 µL RNase inhibitor, all kept on ice. Brain tissue was homogenized with 2–3 strokes, and the resulting cell suspension was passed through a 70 µm cell strainer (Falcon) placed over a 50 mL conical tube on ice. If residual tissue remained, it was re-homogenized with 2–3 additional strokes, and the resulting suspension was similarly filtered. The filter was rinsed twice with 1 mL of dounce buffer, and the remaining cells were pushed through using the plunger of a 1 mL syringe. Filtered cells were aliquoted into 2 mL microcentrifuge tubes and centrifuged for 30 seconds at 10,000 RPM (9.3 RCF) at 4°C. The resulting brain cell pellets were resuspended in a 30% Percoll gradient (GE Healthcare, Princeton, NJ, USA) and centrifuged at 700 × g for 10 minutes. The supernatant containing myelin was discarded, and the cell pellets were washed with Hank’s Balanced Salt Solution (HBSS) containing 2% FBS, followed by immunolabeling for flow cytometry analysis.

### Flow cytometry

Single-cell suspensions were incubated with fluorophore-conjugated antibodies in PBS supplemented with 2% (vol/vol) fetal bovine serum at 4 °C for 15 minutes. Dead cells were excluded using Ghost Dye Violet 510 (Tonbo Biosciences). The following antibodies were used for cell surface immunolabeling: TMEM119 (106-6), CD11b (M1/70), CD86 (GL-1), CD45 (13/2.3), VISTA (MIH63), and TREM2 (6E9). For intracellular staining, brain cells stained with surface Abs were fixed and permeabilized using a fixation/permeabilization buffer, followed by immunolabeling with antibodies against APOE (EPR19392) and BCL11A (14B5), according to the manufacturer’s instructions (Invitrogen). All antibodies were purchased from BioLegend or Abcam unless otherwise specified. Flow cytometry data were acquired using the Attune NxT flow cytometer (Invitrogen) and analyzed using FlowJo software (version 10.6, TreeStar).

## Acknowledgements

We thank AtlasXomics for generating the spatial ATAC-seq data, and the Histology Core at the University of North Dakota (UND) for tissue embedding and cryosectioning. We also acknowledge support from the Phase II COBRE grant and the Genomics Core at UND.

## Author Contributions

X.W. and K.Y. conceived the project and supervised the research. D.K. and H.H. contributed equally to the experimental design and data analysis. A.Z., and L.Li performed chromatin accessibility and spatial-ATAC-seq analyses. R.M. assisted with experimental setup and mouse brain tissue processing. D.D. contributed to initial assessment of section anatomy and cell type specification. M.T., L.Lu, J.P. provided guidance on data interpretation. J.Y. and K.Y. performed microglial isolation and flow cytometry experiments. X.W. led manuscript writing, with contributions from all co-authors.

## Data and materials availability

The raw data reported here is available at BioSample database with accession number SAMN48268985 and SAMN48268986.

## Ethics approval and consent to participate

Not applicable. Studying does not involve human subjects. No ethical approval or consent are required.

## Consent for publication

Not applicable.

## Supplementary Information

**Fig. S1.**
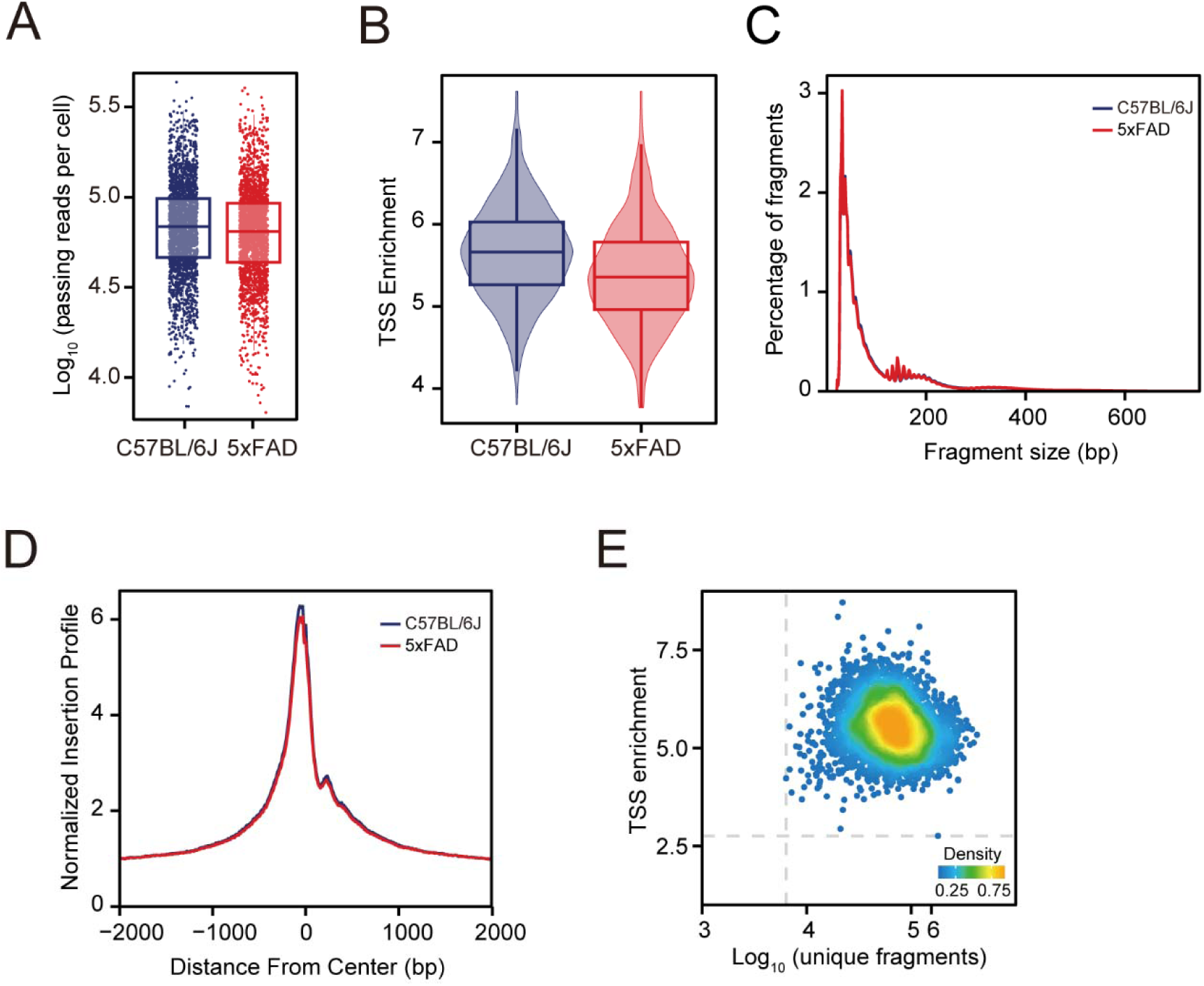
Spatial-ATAC-seq data quality. (A) Comparison of the number of unique fragments between 5×FAD and control. (B) Comparison of the fraction of TSS fragments between 5×FAD and control. (C) Comparison of the insert size distribution of ATAC-seq fragments between 5×FAD and control. (D) Comparison of the enrichment of ATAC-seq reads around TSSs between5×FAD and control. (E) QC filtering plot showing TSS enrichment score versus unique nuclear fragments per pixel. Dot color represents point density. Most data points exhibit high TSS enrichment and fragment counts, indicating high data quality.

**Fig. S2.**
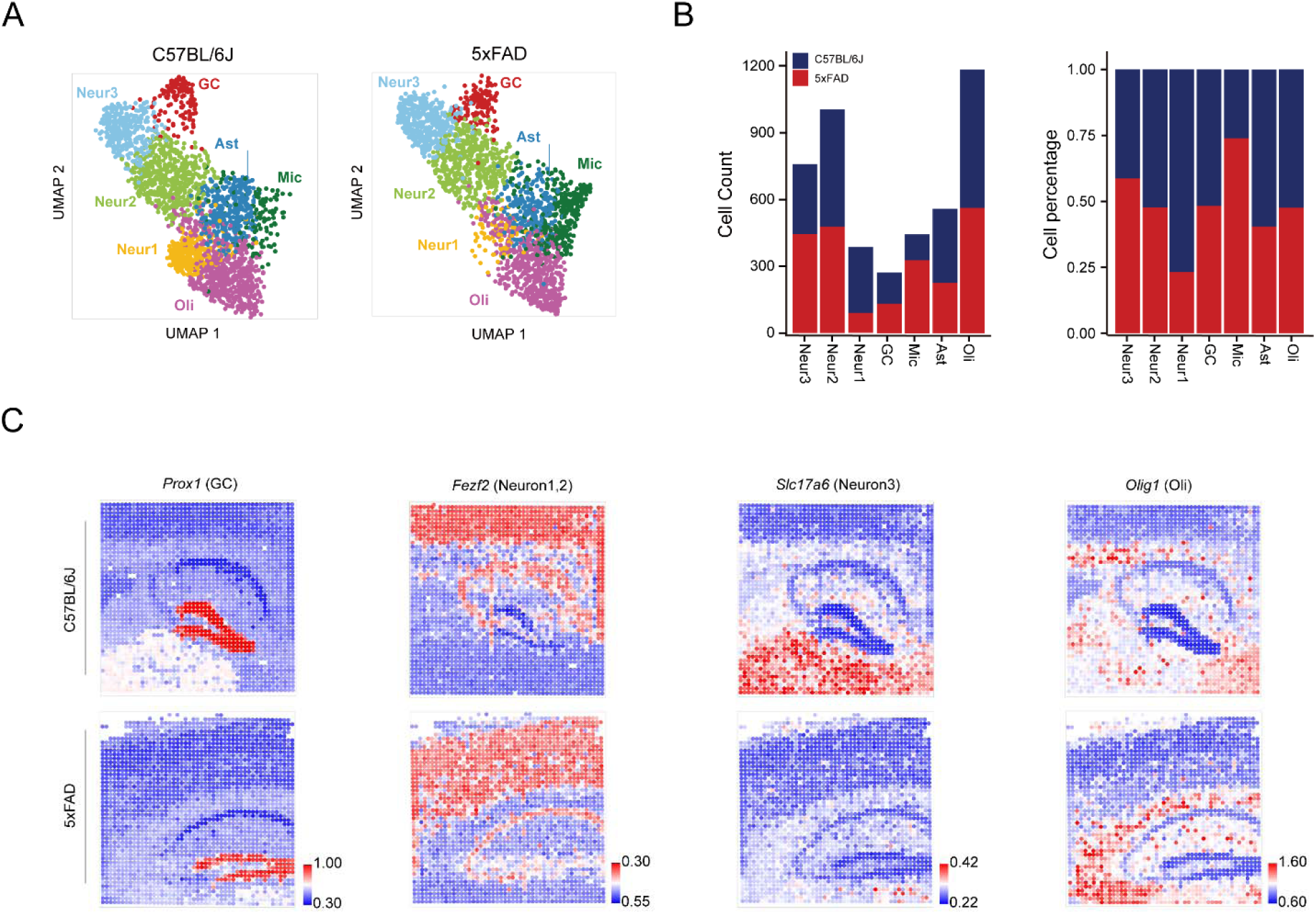
Sample and cluster basic information statistics. (A) UMAP for 5×FAD and control samples. (B) Number of spots (left), percentage of number of spots (right) for 5×FAD and control samples in different clusters. (C) Spatial expression patterns of selected marker genes across seven cell types: *Prox1* (Granule cells), *Fezf2* (Neuron 1, 2), *Slc17a6* (Neuron 3), *Olig1* (oligodendrocytes).

**Fig. S3.**
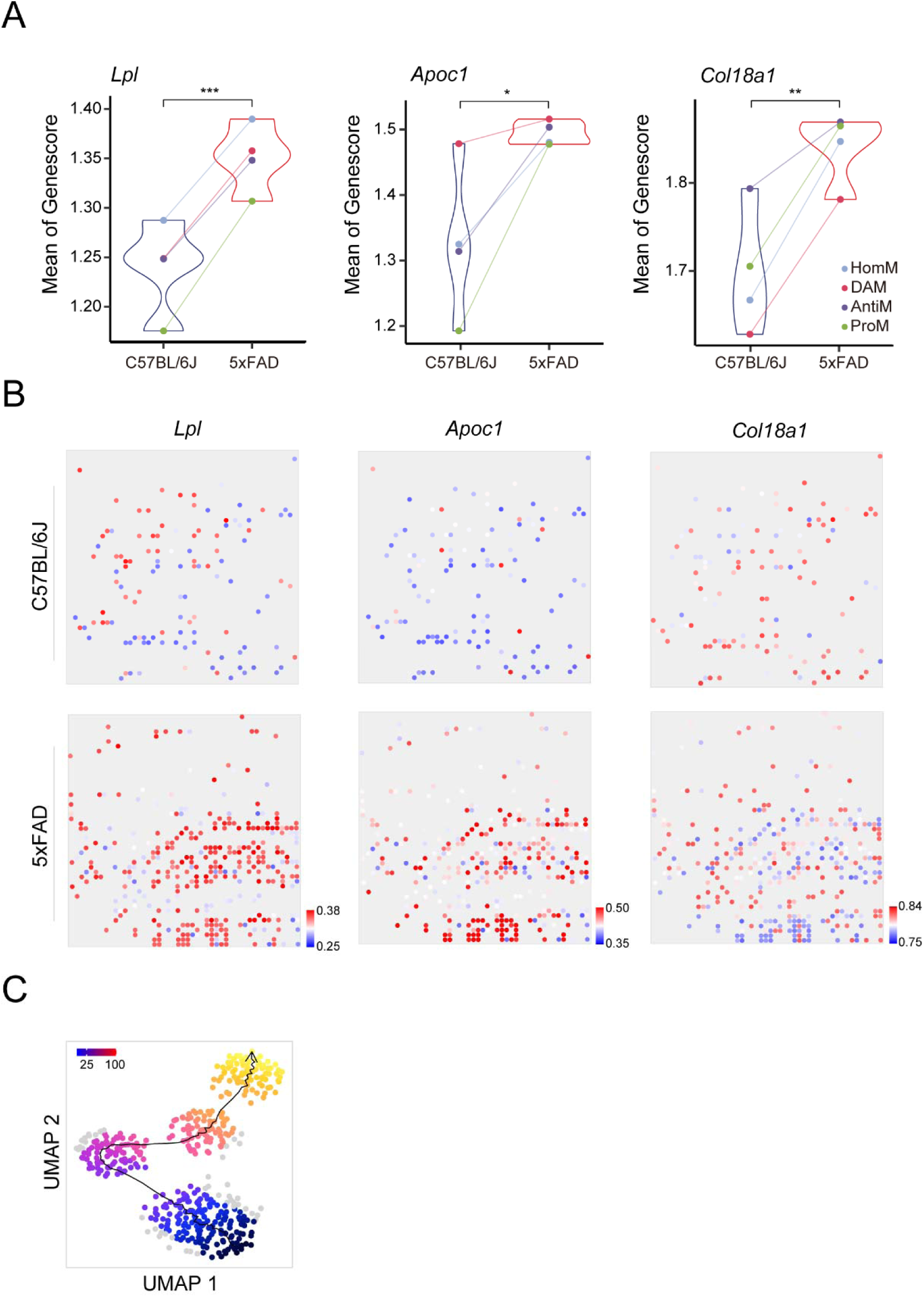
Differences in the expression and distribution of selected genes in microglia from 5×FAD and control. (A) Gene score comparisons for selected genes between 5×FAD and its control, grouped by microglia subtypes. Asterisks denote statistically significant differences. (B) Spatial gene expression profiles for selected genes in C57BL/6J (top) and 5×FAD (bottom) mouse models. Colors represent expression levels. (C) Pseudo-time trajectory analysis showing transitions between microglia subtypes.

**Fig. S4.**
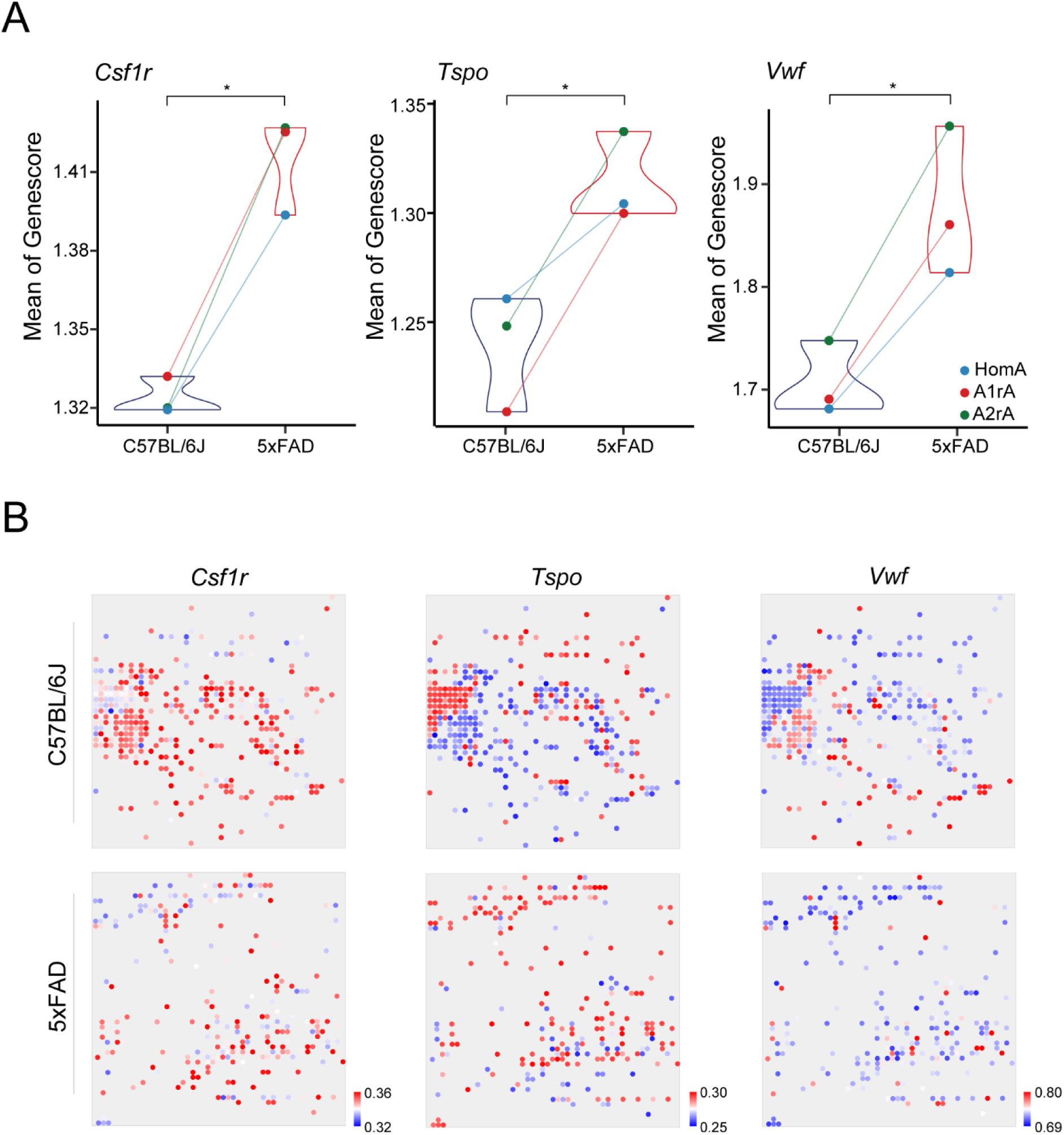
Differences in the expression and distribution of selected genes in astrocyte from 5×FAD and control. (A) Gene score comparisons for selected genes between 5×FAD and its control, grouped by astrocyte subtypes. Asterisks denote statistically significant differences. (B) Spatial gene expression profiles for selected genes in C57BL/6J (top) and 5×FAD (bottom) mouse models. Colors represent expression levels.

**Fig. S5.**
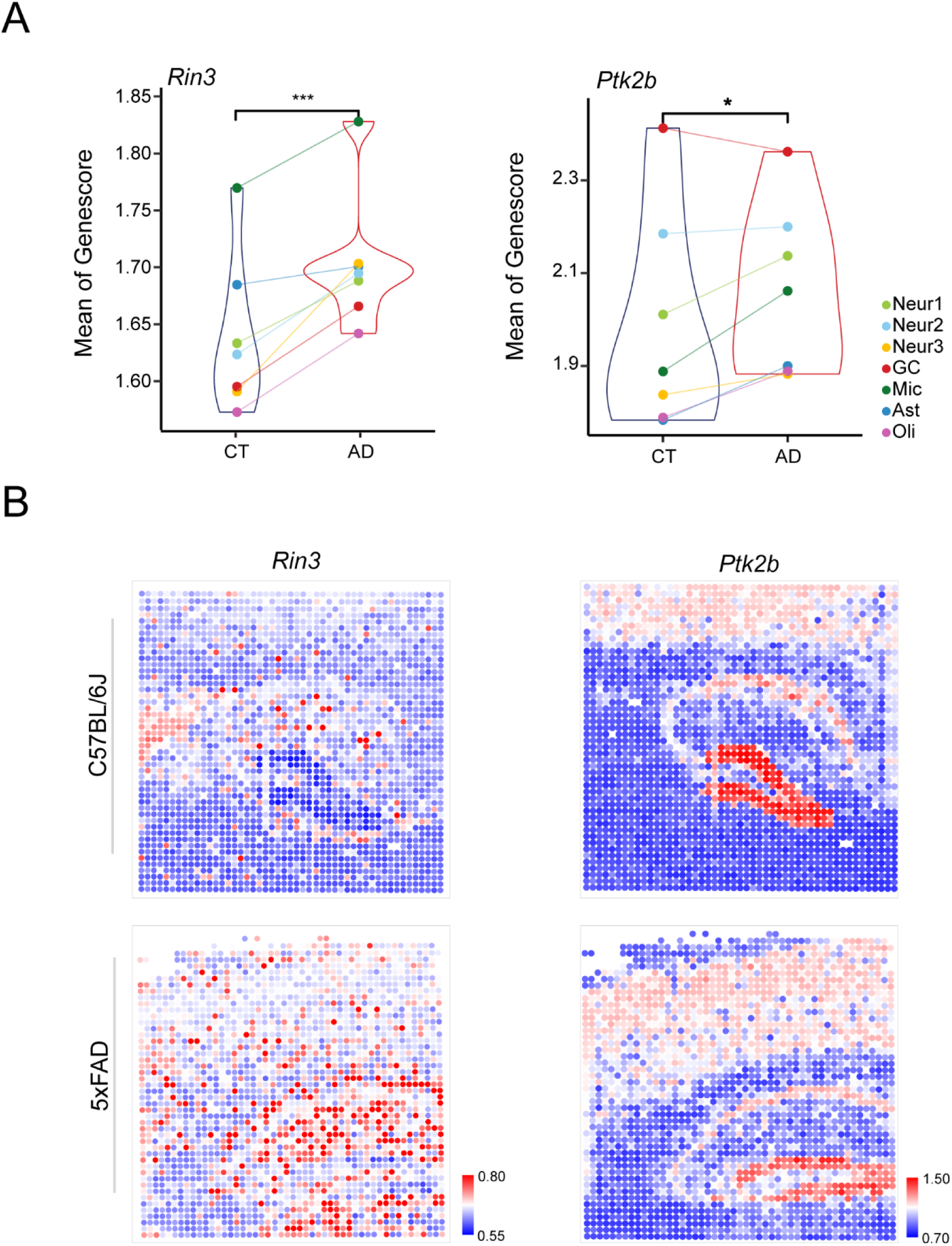
Differences in the expression and distribution of *Bcl11a* downstream genes from 5×FAD and control. (A) Gene score comparisons for selected genes between 5×FAD and its control, grouped by cell types. Asterisks denote statistically significant differences. (B) Spatial gene expression profiles for selected genes in C57BL/6J (top) and 5×FAD (bottom) mouse models. Colors represent expression levels.

**Fig. S6.**
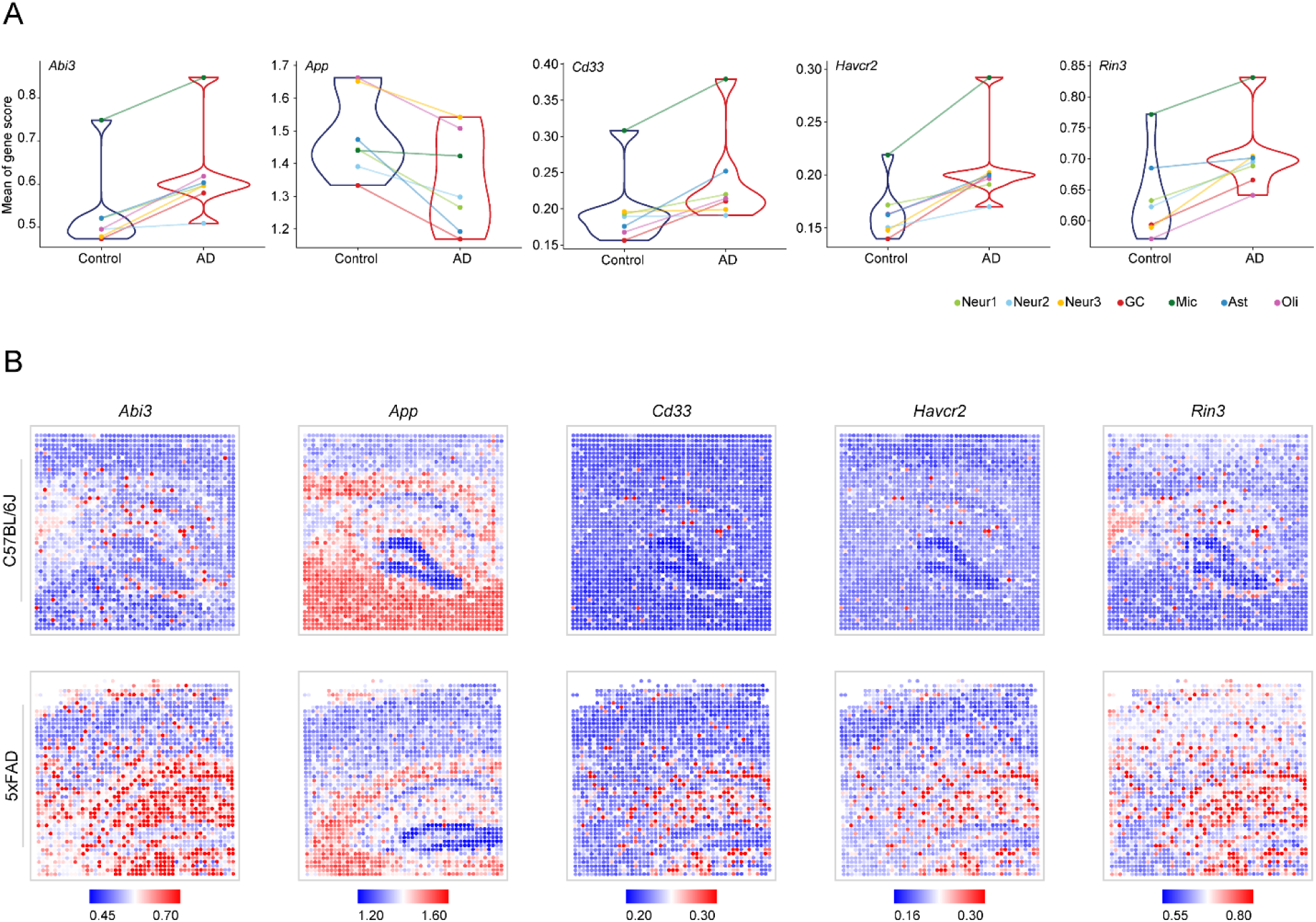
Differences in the expression and distribution of AD risk genes with differential expression from 5×FAD and control mice. (A) Gene score comparisons for selected genes between 5×FAD and C57BL/6J control mice, grouped by cell types. Asterisks denote statistically significant differences. (B) Spatial gene expression profiles for selected genes in C57BL/6J (top) and 5×FAD (bottom) mouse models. Colors represent expression levels.

